# Pat1 globally activates deadenylation-dependent decapping to clear inefficiently translated oligo(A)-tailed transcripts

**DOI:** 10.64898/2026.10.22.760927

**Authors:** Lucie Labeauvie, Jackson Peter, Hélène Zuber, Bertrand Séraphin

## Abstract

In eukaryotes, mRNA turnover occurs mainly through the 5’ - 3’ mRNA decay pathway and involves deadenylation, decapping, and exonucleolytic mRNA body digestion. While the conserved protein Pat1 has initially been described as an activator of decapping for some deadenylated mRNAs, the role of deadenylation-dependent decapping has recently been questioned. Hence, Pat1 has alternatively been suggested to regulate specific mRNA decay pathways, given its modest impact on overall transcriptome composition. Here, we combine multi-omic and genetic approaches to characterize a *PAT1* deletion mutant in *Saccharomyces cerevisiae*. Although transcriptome and proteome analyses confirm that only a limited subset of genes is affected by Pat1 inactivation, poly(A)-tail sizing demonstrates that Pat1 loss leads to transcriptome-wide accumulation of capped oligo(A)-tailed mRNAs. Together with the observation that oligo(A)-tailed mRNAs are bound by Pat1 *in vivo* in wild-type cells, this indicates that Pat1 is a general activator of deadenylation-dependent decapping. We further show that oligo-adenylated mRNAs are translated in *Δpat1* cells, albeit inefficiently. Altogether, our data indicate that Pat1 mediates the clearance of oligo(A)-tailed mRNAs protecting cells from the ensuing inefficient translation, without inducing major changes in overall transcript distribution.

## Introduction

Eukaryotic mRNAs are central actors of gene expression; thus, their levels are tightly controlled to adapt protein production to cellular demands. This involves several processes, including cytoplasmic mRNA turnover that is critical to eliminate unwanted transcripts and modulate protein production. Among the two major mRNA decay pathways, the 5’-3’ pathway plays a dominant role in degrading worn transcripts^1–3^. This process involves three essentially sequential steps: (i) shortening of the poly(A) tail, (ii) removal of the 5’ m^7^G cap, and (iii) Xrn1- mediated 5’-3’ exonucleolytic degradation of the deprotected mRNA^4–6^. Early studies on selected mRNAs reported that deadenylation kinetics are rate-limiting for decay, with stable mRNAs undergoing slow deadenylation and short-lived transcripts exhibiting rapid poly(A) tail trimming^7^. This model was further supported by the characterization of two assemblies with deadenylase activities: Pan2/Pan3 and CCR4-NOT complex. The latter is the primary cellular deadenylase and its activity is regulated by a diverse array of activators and inhibitors of mRNA decay^8^.

Early studies suggested a molecular mechanism to target deadenylated mRNAs for decapping. It involves the recognition of residual oligo(A) tails by Lsm proteins, which recruit the decapping complex via the scaffolding protein Pat1^9,10^. This proposal was supported by the observation that mRNA decay rates were reduced in the absence of Pat1^9–11^, even though this was only observed for some unstable transcripts. More recently, the role of deadenylation-dependent decapping in yeast has been challenged by transcriptome-wide analyses. Using Nanopore sequencing to assess poly(A) tail lengths, a study confirmed that deadenylation rates vary in a transcript-specific and codon-usage-dependent manner in optimal growth conditions. But these authors also reported that deadenylation is not a pre- requisite for decapping for certain mRNAs and/or in certain conditions, suggesting that the link between poly(A) tail shortening and cap removal is more complex than previously described^12^. Hence, transcripts encoding ribosomal proteins were proposed to undergo degradation in a nuclear export-dependent but deadenylation-independent manner under heat stress. A parallel analysis, also based on Nanopore sequencing, concluded that decapping rates primarily control mRNA decay in yeast, independently of initial deadenylation rates^13^. The role of deadenylation in controlling mRNA decay and decapping is thus now-a-day questioned.

Additional insights into mRNA decay have emerged from studies of the decapping enzyme itself. The Dcp2 enzyme, in complex with its obligate partner Dcp1 (and Edc4 in mammals), is the primary entity responsible for m^7^G cap elimination^6,14–17^. Genetic and biochemical analyses have revealed that the low intrinsic activity of Dcp2 is tightly regulated by a network of decapping activators—including Edc1, Edc2, Edc3, Scd6, Lsm1, Pat1, Pby1, Dhh1 and Upf1 in yeast and their homologs in other species. These factors ensure that only bona fide targets are irreversibly decapped^6^. Structural and biochemical studies have elucidated how these factors interact with Dcp2 to facilitate its recruitment and/or activation^18–22^. To identify mRNA populations regulated by these activators, transcriptome analyses have been performed in strains lacking individual decapping activators. Hence, Upf1 was shown to be required for the degradation of transcripts containing a premature stop codon without prior deadenylation^23,24^. More recently, it was proposed that Upf1 is also involved in the degradation of mRNAs harboring short poly(A)-tailed independently of the presence of a premature termination codon^25^. For the other decapping co-activators, studies which compared mRNA levels between wild-type and mutant strains grown at optimal temperatures, revealed overall modest transcriptome changes, with only a few mRNAs accumulating in individual mutants. These findings were interpreted as evidence that each decapping activator controls a specific and limited mRNA population, with some transcripts subject to redundant regulation by several factors^26–29^. This interpretation aligns with the observation that inactivation of single decapping activators typically results in mild growth phenotypes, which are exacerbated in strains carrying multiple mutations. Yet, these analyses failed to explain why deletion of the *PAT1* gene (or the gene encoding its partner Lsm1) results in more severe growth phenotypes than those generated by inactivation of other decapping activators. Indeed, elimination of Pat1 generates a severe growth defect in optimal growth conditions that is exacerbated at high temperature^9–11,30^.

Altogether, the precise role of Pat1 remains controversial. While most studies link Pat1 to decapping and mRNA decay, its function appears multifaceted. For instance, Pat1 has been proposed to stabilize autophagy-related transcripts from 3’-5’ degradation during nitrogen starvation-induced autophagy^31^, yet it has inversely been suggested to lower the levels of transcripts involved in autophagy in rich media^29^. Similarly, Pat1 has been implicated in both the repression^29,32^ and activation^30^ of translation, globally or for specific mRNAs, highlighting possibly complex and context-dependent regulatory roles.

In this study, we use multi-omic analyses to re-investigate mRNA decay and the role of Pat1 in this process in the yeast *S. cerevisiae*. Our analyses uncover a transcriptome-wide perturbation in *Δpat1*, undetectable with classical transcriptomic or proteomic analyses, revealing that the absence of Pat1 results in a widespread accumulation of oligo-adenylated mRNAs. Our data confirm that the absence of Pat1 results in a decapping defect and support the existence of a general deadenylation-dependent decapping pathway. We observe translation of oligo-adenylated mRNAs accumulating in the Δ*pat1* strain and concomitantly overall protein production is slowed-down. Using reporters, we show further that short oligo(A) tails reduce translation efficiency and that this defect is exacerbated under suboptimal growth conditions in both wild-type and *Δpat1* cells. Overall, our data confirm the implication of Pat1 in deadenylation-dependent decapping and reveals its role in the sanitization of oligo(A)-tailed transcripts which are poorly translated.

## Results

### Limited transcriptomic and proteomic alterations in a Δpat1 mutant despite severe growth defects

The observation that Δ*pat1* strains exhibit a significant growth defect compared to wild-type (WT) strains across all tested temperatures (Fig. 1A) suggests a central molecular dysfunction in these cells. This phenotype is exacerbated at elevated temperatures, with no growth observed at 37°C (Fig. 1A). However, recent transcriptomic analyses have reported only modest changes in mRNA abundance in a Δ*pat1* strain^27,29^, a finding that appears counterintuitive given the marked growth impairment. This discrepancy is compounded by the observation that Δpat1 and Δdhh1 share similar transcriptome alterations^29^, despite these mutants having distinct growth phenotypes, and their combination exacerbating the Pat1 poor growth^45^.

**Figure 1.**
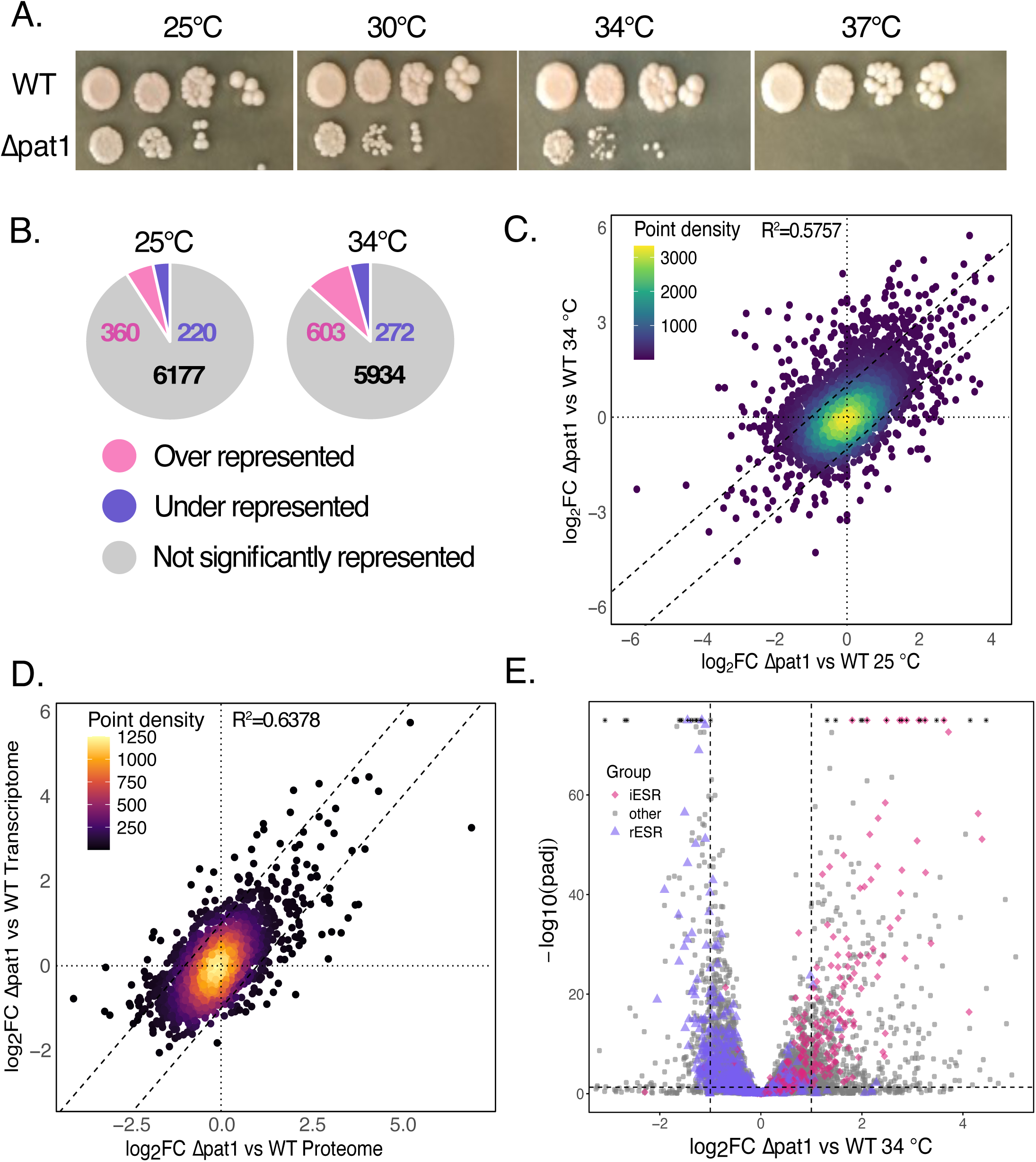
The absence of Pat1 has only a minor quantitative impact on the transcriptome and proteome, despite inducing strong phenotypic effects. (A) The deletion of *PAT1* leads to a thermosensitive phenotype. Yeast growth assays were performed using the WT and Δ*pat1* strain. Serial 10-fold dilution of cultures grown at 25°C were plated drop-wise on YPDA-agar media and incubated at the indicated temperatures for 2 days. (B) Differentially represented transcripts between the WT and Δ*pat1* strain account for a minor transcriptome portion. The number of significantly over- (pink) and under- (violet) represented or non-significantly affected (grey) transcripts are represented as a pie chart for Illumina-based transcriptome sequencing at 25 and 34°C. (C) Differentially affected transcripts between the WT and Δ*pat1* strain grown either at 25°C or 34°C are highly correlated with increased differences at the near-restrictive temperature. The log2 Fold Change of the comparison of Δ*pat1* and WT at 25°C and 34°C is compared and represented as a density plot for commonly found genes (n = 6727, ribosomal RNAs are excluded). The Spearman correlation of the two dataset is represented as R^2^. (D) The proteome reflects the transcriptome. The log2 Fold Change of the comparison between the WT and Δ*pat1* strain at the transcriptomic and proteomic level at 34°C was compared and represented as a density plot for commonly found features (n = 2672). The Spearman correlation of the two dataset is represented as R^2^. (E) The Δ*pat1* transcriptome reflects a stress-induced situation overlapping a reduced mRNA decay. Volcano plot representation of the differentially represented transcripts between the WT and Δ*pat1* strain at 34°C, with induced and repressed environmental stress response genes (pink diamonds and violet triangles, respectively) highlighted. Transcripts with a −log_10_(p-adj) greater than 75 are shown with black crosses and capped at 75 to improve readability. The horizontal dashed straight lines indicate p-value of 0.05 and vertical dashed straight lines log2FC= +/-1.

To investigate this apparent paradox, we performed transcriptomic and proteomic analyses of WT and Δ*pat1* cells at both permissive (25°C) and near-restrictive (34°C) temperatures. Consistent with previous studies^27,29^, using short read sequencing, we observed limited transcriptomic changes in Δ*pat1* at 25°C: only 580 RNAs out of 6,177 exhibited a fold change greater than 2 (log₂FC > 1) with a false discovery rate (FDR) of 0.05 (p-adjusted < 0.05), comprising 360 up-regulated and 220 down-regulated transcripts (Fig. 1B). At 34°C, the number of differentially expressed genes increased only modestly to 875 (603 up-regulated and 272 down-regulated, Fig. 1B). A comparison of the transcriptomic changes between the two temperatures revealed a good correlation, with only a small subset of genes showing a modest exacerbation of differential expression at 34°C (Fig. 1C).

Given that phenotypic consequences are ultimately mostly mediated by the proteome^46^, we performed label-free quantitative proteomic analysis using mass spectrometry on WT and Δ*pat1* samples prepared under the same near-restrictive conditions used for the transcriptomic analyses. Despite the lower depth of proteomic coverage compared to transcriptomics (products of 2843 versus 6809 genes detected, respectively), we observed a similar pattern of changes with the limited transcriptomic differences observed in Δ*pat1* being mirrored at the proteomic level (Fig. 1D). Thus, neither a striking transcriptome alteration nor unbalanced translation of different mRNAs explains the strong Δ*pat1* phenotype.

Gene Ontology (GO) enrichment analysis of up-regulated transcripts in the Δ*pat1* strain at 34°C revealed an overrepresentation of metabolic and biosynthetic pathways involved in energy production and nutrient processing. In contrast, down-regulated transcripts were strongly associated with cytoplasmic translation including ribosome biogenesis, consistent with the reduced growth of the mutant (Fig. S1). Notably, the Environmental Stress Response (ESR) in yeast is a coordinated transcriptional program that reallocates resources from growth to survival^47^. The induced ESR (iESR) globally promotes cell protection while the repressed ESR (rESR) globally downregulates growth-related processes. Globally, iESR genes^48^ were up- regulated in the Δ*pat1* at 34°C while rESR genes were generally down-regulated (compare the distribution of transcripts respectively indicated in pink and violet in Fig. 1E). However, this effect was asymmetrical: 50% of known iESR genes^48^ were significantly up-regulated in the Δ*pat1* at 34°C whereas only 13% of rESR genes were significantly down-regulated (Fig. 1E). This uneven effect reflects the fact that Pat1 deletion disproportionately affects the transcriptome, with a greater number of up-regulated transcripts exhibiting stronger fold changes, while fewer down-regulated transcripts are more modestly affected (Fig. 1E). Consequently, the Δ*pat1* transcriptome likely reflects both a combination of an ESR response overlapping with a global mRNA decay defect, the latter stabilizing mRNAs, thereby amplifying the effects on induced transcripts and buffering the repression of others.

In summary, even under near-restrictive conditions, Δ*pat1* exhibits limited quantitative changes in transcript and protein contents compared to the wild-type control, a finding that contrasts sharply with its severe growth defect. The observed variations in the levels of mRNAs are consistent with a global stress response, superimposed on widespread transcript stabilization due to the loss of a mRNA decay factor. Thus, the most affected mRNAs are unlikely to be direct targets of Pat1, in contrast to previous suggestions^27,29^.

### Pat1 inactivation globally shortens poly(A)-tail lengths across the transcriptome

Given that short-read analyses failed to reveal a strong quantitative impact of Pat1 on mRNA levels, we performed long-read sequencing, which enables both qualitative and quantitative transcriptome monitoring. Since poly(A)-tail length is closely associated with mRNA translation and decay, we conducted a FLEP-Seq2 analysis^38^ on both wild-type and *Δpat1* strains to specifically assess poly(A)-tail status in these genetic backgrounds.

Quantitatively, the FLEP-Seq2 results were largely consistent with short-read sequencing, with no notable strain-specific deviations (Fig. S2). Furthermore, no differences in splicing were observed between the WT and *Δpat1* strains as proportion of retained-introns containing mRNAs was not significantly different (p-value=0.1, Wilcoxon test). However, this analysis revealed a striking difference in mRNA poly(A)-tail lengths between the two strains. We essentially detected poly(A)-tail lengths ranging from 5 to 100 adenines, consistent with the maximum sizes estimated by other methods^12,49,50^. When we analyzed the distribution of poly(A)-tail lengths for transcripts detected at least 25 times in each condition and shared between the WT and *Δpat1* strains, a clear size difference emerged (Fig. 2A). In the WT strain, the proportion of mRNAs was highest for transcripts with poly(A)-tails of at least 20 adenines, with a gradual decline in proportion for tails up to 70 adenines (median size value for all reads: 32). In stark contrast, *Δpat1* exhibited a poly(A)-tail length profile that peaked at 10 adenines, followed by a sharp decrease up to 25 adenines and a more gradual decline up to 70 adenines (median size value for all reads: 20). These results demonstrate that the majority of the *Δpat1* transcriptome consists of mRNAs with significantly shorter poly(A)-tails compared to WT.

**Figure 2.**
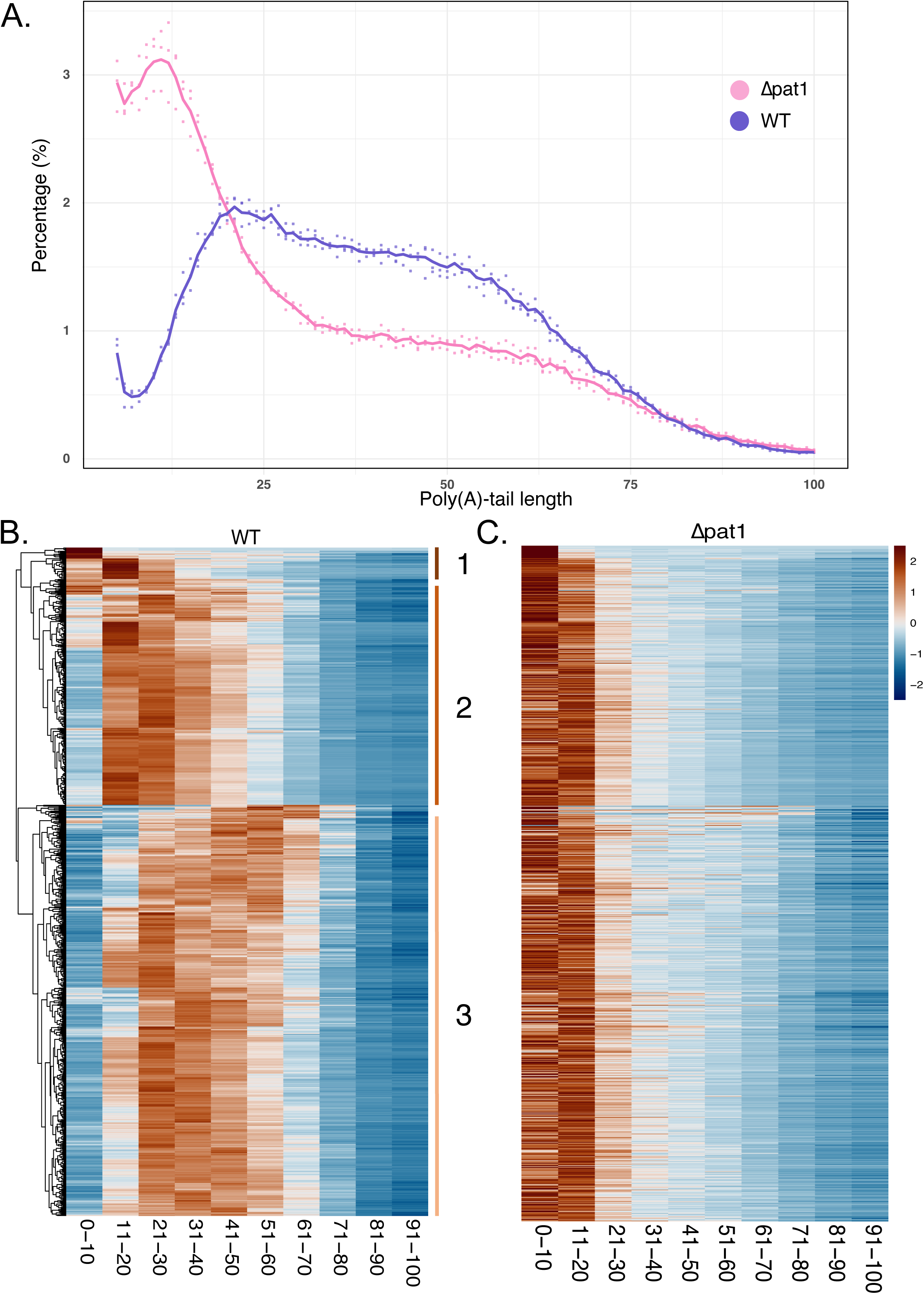
The absence of Pat1 leads to the accumulation of short-tailed mRNA for all transcripts. (A) Weighted poly(A)-tail distribution for the WT (violet) and Δ*pat1* strain (pink) from 5 to 100A with individual points as biological samples and average by genotypes as a straight colored line for 2951 genes. (B) Heatmap showing the poly(A) tail lengths by bin of 10 nucleotides for all mRNAs in the wild-type strain. The percentage of reads according to their poly(A)-tail length was calculated accounting all transcripts present at least 25 times and with a poly(A)-tail length capped at 100A. Percentages were z-score rescaled. Heatmaps were ordered based on the poly(A)-tail similarity profiles. Three transcript clusters are indicated on the right. The scale is provided in Panel C. (C) Heatmap showing the accumulation of short-tailed transcripts for all mRNA in the Δ*pat1* strain. The data were processed as for panel B and mRNAs are ordered according to the order used in panel B.

To determine whether this phenomenon was global or restricted to specific mRNAs, we calculated the percentage of transcripts with poly(A)-tails grouped in 10-adenine increments (from 0 to 100 adenines) for each mRNA. The results are presented as a heatmap, clustered according to the poly(A) profile in the WT strain (Fig. 2B, left). Distinct groups of transcripts became visible, consistent with previous analyses^12,51,52^. In particular, the ribosomal protein genes (RPG) cluster had overall shorter poly(A)-tail lengths compared to non-RPG transcripts (compare cluster 2 and 3; Fig. 2B). The abnormally short-tailed transcripts in the minor cluster 1 corresponds to transcripts of retro-transposable elements and dubious open reading frames, likely being destabilized faster than other RNAs.

When we plotted the poly(A)-tail size of mRNAs in the Δ*pat1* strain in the same order, no obvious cluster emerged, indicating a total loss of the regulation of poly(A)-tail lengths. We observed that ∼90% of all mRNAs in Δpat1 had the majority of their transcripts with a poly(A)- tail ranging from 0-10 or 11-20 adenines, confirming that the distribution in Figure 2A was not skewed by a specific mRNA population (Fig. 2C). Clustering according to the poly(A) profile in Δ*pat1* (Fig. S3) shows mRNAs with a predominant transcript proportion between 11-20 adenines (Cluster A, 1 020 739 mRNAs), with others comprised between 0 and 10 residues (Cluster B, 330 112 mRNAs). As the poly(A) tail length of transcripts in Δ*pat1* sharply peaks at 11 nucleotides (Fig.2A), this artificially divides the distribution based on incremental groups of 10A. Indeed, GO analyses failed to identify significant features specifically associated with transcripts of clusters A and B (Fig. S4), suggesting that the difference between these two populations had no functional basis but rather resulted from variations around the poly(A) tail length peak value.

Taken together, while classical RNA-Seq failed to detect widespread changes in mRNA levels in *Δpat1*, our FLEP-Seq2 analysis reveals that *PAT1* deletion globally disrupts poly(A)-tail homeostasis, leading to accumulation of oligo(A)-tailed mRNAs. This effect is not restricted to specific mRNA subsets but instead represents a general defect.

### Inefficient decapping leads to accumulation of oligo(A)-tailed mRNAs in the absence of Pat1 and limits growth

To further characterize the mRNA population in the *Δpat1* strain, we investigated whether the oligo(A)-tailed mRNAs that accumulate in this mutant are still capped. In wild-type cells, mRNAs are essentially capped with only a minor fraction of decapped mRNA decay intermediates present. Thus, mRNAs extracted from wild-type cells are essentially not susceptible to *in vitro* degradation by Xrn1, unless they are first decapped as shown for the *ACT1*, *RPL41B* and *PGK1* mRNAs in Fig. 3A. Analyses of mRNAs extracted from Δ*pat1* cells indicate that they are similarly protected by a cap from Xrn1 degradation (Fig. 3A). Because the transcriptome of Δ*pat1* cells is essentially constituted from oligo-adenylated mRNAs, this observation argues that a large fraction of the transcripts is capped. To assess directly the presence of an m^7^G cap on oligo-adenylated mRNAs present in the Δ*pat1* strain, we immunoprecipitated mRNAs using an anti-m^7^G cap antibody and analyzed the poly(A)-tail lengths of two representative transcripts, RPL41B and RPL10, using the RACE-PAT technique^41^. RPL41B was selected because its poly(A)-tail length profile (Fig. S5) closely matches that of the global transcriptome (Fig. 2A), while RPL10 exhibited a slightly distinct but still representative pattern (Fig. S5). Although some non-specific precipitation was detected with the negative control, the signal was significantly stronger with the anti-m^7^G cap antibody, confirming enrichment for capped mRNAs (Fig. 3B). Analysis of the poly(A) tails of RPL41B and RPL10 in the immunoprecipitated fractions revealed that the oligo(A)-tailed mRNAs accumulating in the *Δpat1* strain are indeed capped (Fig. 3B).

**Figure 3.**
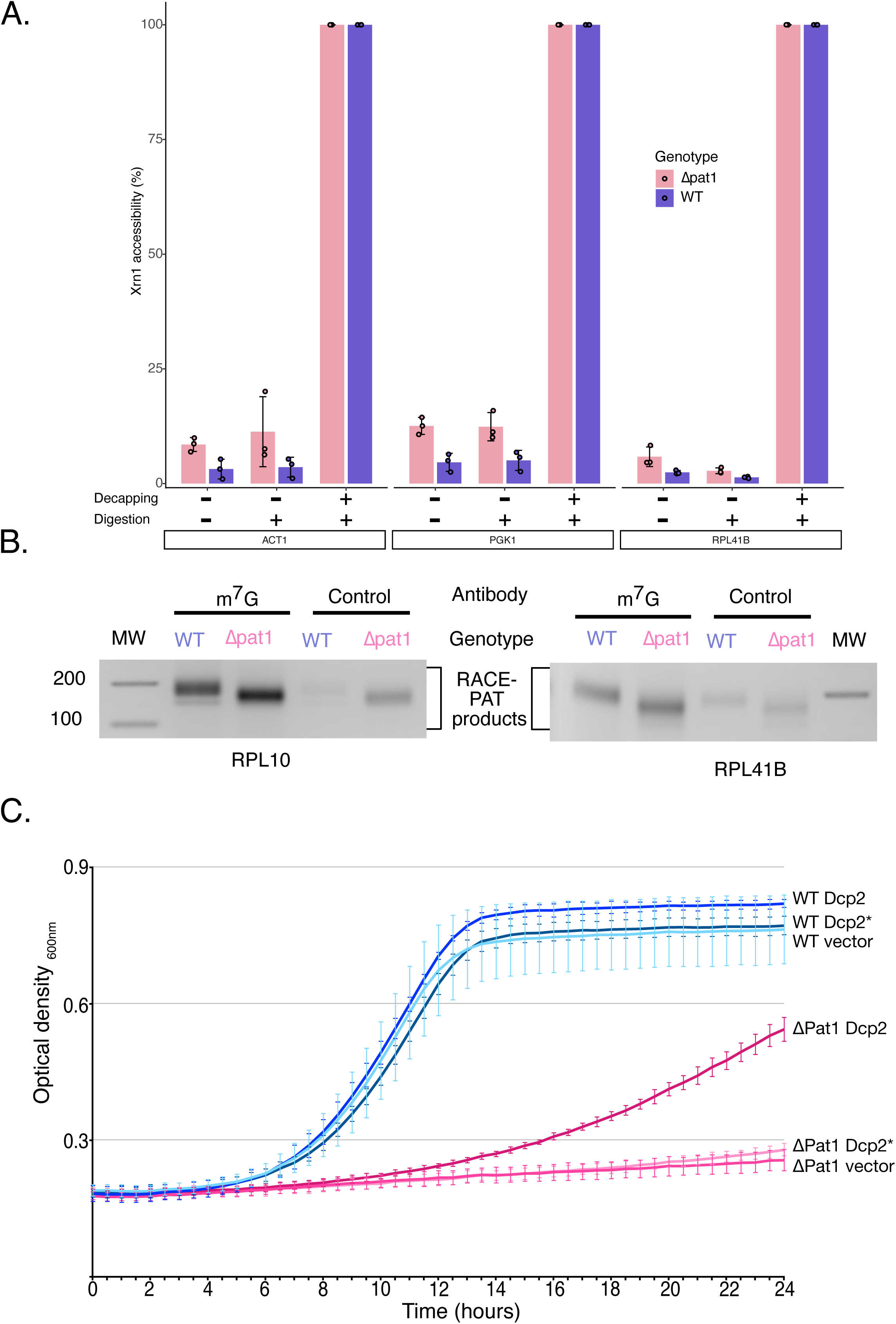
There is a general decapping defect when Pat1 is absent. (A) mRNAs present in Δpat1 strain have a protected 5’ end. The presence of caps protecting mRNAs encoding different proteins in wild-type (violet) and Δ*pat1* (pink) strain was assess through their susceptibility to Xrn1 digestion *in vitro* and monitored by RT-qPCR. Maximal susceptibility was determined by treating total RNA samples with Xrn1 after enzymatic cap removal. Biological triplicates were analyzed (n=3) and technical triplicates were performed for each RT-qPCR reaction. (B)Short-tailed mRNAs are still capped in the Δ*pat1* strain. RACE-PAT analysis of the RPL10 and RPL41B mRNAs from the wild-type or Δ*pat1* strains after precipitation with anti-m7G or control antibodies were made in biological duplicates. Pictures of ethidium bromide-stained 3% agarose gels are presented. MW= molecular weight. (C) Overexpression of the decapping holoenzyme (Dcp2/Dcp1) suppresses the growth phenotype of the Δ*pat1* strain. Yeast growth assays in liquid YPDA medium were performed using the WT and Δ*pat1* strain after transformation with plasmids overexpressing Dcp2 full length and Dcp1 (termed Dcp2/Dcp1), catalytically inactive (E152A E153A) Dcp2 full length and Dcp1 (termed Dcp2*/Dcp1), or plasmids without inserts. The OD_600nm_ was monitored every 30 minutes over a 24-hour period. Standard deviations of biological triplicates (n=3) are shown as straight vertical lines.

The accumulation of capped oligo(A)-tailed mRNAs in the absence of Pat1 suggests that inefficient decapping may be limiting growth in Δ*pat1*. To test this hypothesis, we examined whether overexpression of the decapping enzyme could rescue the growth defect observed in the mutant. Wild-type and *Δpat1* cells were transformed with plasmids expressing the two subunits of the decapping holoenzyme, Dcp1 and Dcp2, under the control of strong constitutive promoters. A catalytically inactive mutant of Dcp2^19^ (Dcp2 E152A E153A, hereafter referred to as Dcp2*) and an empty vector were used as negative controls. Overexpression of the Dcp2/Dcp1 holoenzyme suppressed the slow-growth phenotype of *Δpat1* cells compared to those transformed with the empty vector. As often in such assays, suppression was partial because increased protein levels could not fully compensate for absence of efficient decapping activation mechanisms. Importantly, this suppression was dependent on the catalytic activity of Dcp2, as no rescue was observed with Dcp2*. None of the constructs significantly affected the growth of wild-type cells, indicating that their expression did not induce toxic effects (Fig. 3C).

Together, our biochemical and functional analyses demonstrate that decapping is impaired in the absence of Pat1, leading to the accumulation of capped oligo(A)-tailed mRNAs and likely contributing to the growth defect observed in *Δpat1*.

### Pat1 associates with transcriptome-wide oligo(A)-tailed mRNAs in a wild-type strain

The transcriptome-wide accumulation of oligo(A)-tailed mRNAs in the Δ*pat1* strain, and the associated decapping defect, could theoretically arise as an indirect consequence of the mutation. Given that the Lsm1-7 complex—known to associate with Pat1^9–11,45,53–56^—has been reported to interact with oligo(A)-tailed *in vitro* mRNAs^57,58^, we sought to determine whether Pat1 itself associates with oligo(A)-tailed mRNAs *in vivo* and to establish simultaneously the presence of these species in wild-type Pat1 context. This question is particularly relevant, as some previous studies have downplayed the role of decapping promoted by deadenylation.

To identify mRNAs bound by Pat1 in a wild-type context, we immunoprecipitated TAP-tagged Pat1, recovered the associated mRNAs, and analyzed them using FLEP-Seq2. As a control, we used Pab1-TAP-bound mRNAs. Indeed, Pab1 was reported to preferentially bind mRNAs with poly(A) tails of at least 25 adenosines, although it can also associate with shorter oligo(A) tails^51,59,60^ (Fig. 4A). To ensure that the TAP tags did not alter protein function, we first verified that the growth phenotypes of Pat1-TAP and Pab1-TAP strains were comparable to those of the WT strain and did not mimic the phenotypes observed in the corresponding deletion mutants (Fig. S6).

**Figure 4.**
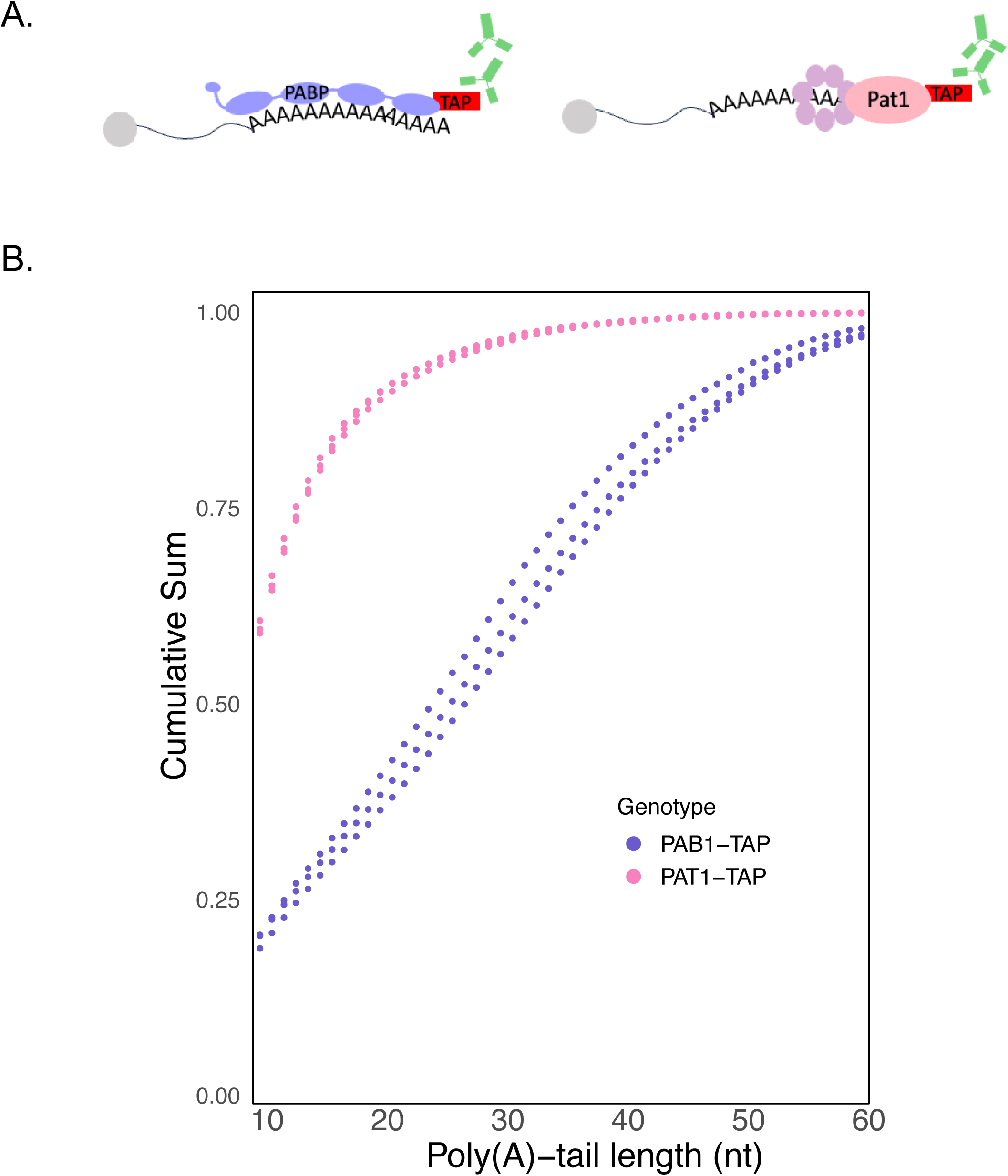
The decapping cofactor Pat1 binds oligo-adenylated mRNAs. (A) Scheme of the RNA-Immunoprecipitation by targeting the TAP-TAG of the protein of interests using as control Peroxidase Anti Peroxidase (PAP). (B) The proportion of transcripts with a tail length shorter than or equal to that value is presented as function of poly(A) - tail length for mRNAs precipitated by Pat1 (pink) or Pab1 (violet). Dots indicate the values obtained from biological triplicates.

FLEP-Seq2 analyses revealed that nuclear-encoded mRNAs associated with Pat1 had shorter poly(A) tails than for Pab1-TAP-bound transcripts. The cumulative distribution of poly(A) tail lengths showed that more than 50% of Pat1-bound mRNAs had fewer than 15 adenines, and 75% had fewer than 20 adenines. In contrast, 50% of Pab1-bound mRNAs had at least 30 adenines (Fig. 4B). Together, these results demonstrate that Pat1 predominantly associates with mRNAs containing oligo(A) tails in a WT context. We observed among the oligo(A)-tailed mRNAs bound to Pat1, some, such as those encoding ribosomal proteins, that have been described in previous publications^13^ as being decapped without prior deadenylation (Fig. S7). Our observations thus contradict these prior conclusions.

To characterize the features of Pat1-bound mRNAs, we analyzed the natural poly(A) tail status of this population in wild-type cells by determining the distribution of their median poly(A) tail length based on our FLEP-Seq2 data. This distribution was then compared to the poly(A) tail size distribution of all mRNAs present in wild-type cells, to assess whether Pat1- bound transcripts are unusual (Fig. S8). We observed no significant difference between the two distributions, indicating that Pat1-bound mRNAs are not intrinsically biased toward a specific median poly(A) tail length but instead reflect the overall transcriptome (Fig. S8).

Overall, our data argue that Pat1 acts as a general decapping coactivator without defined transcript specificity.

### Accumulation of oligo(A)-tailed mRNA in the Δpat1 context leads to an inefficient translation

Poly(A) tails on mRNA are known to stimulate protein synthesis by recruiting the Pab1 protein, which in turn interacts with the eIF4G translation initiation factor^61,62^. Given the accumulation of oligo(A)-tailed mRNAs in Δ*pat1*, we investigated whether this compromised protein production.

To assess protein synthesis in Δ*pat1*, we first quantified translation efficiency using the methionine analog HPG (L-homopropargylglycine). Wild-type and Δ*pat1* cells were grown at 25°C and 34°C in methionine-depleted media, and HPG was added to label newly synthesized proteins. Samples were collected 10 minutes after HPG induction, and cells were fixed, permeabilized, and subjected to a click reaction to label HPG-incorporated proteins with Alexa488^42^. Protein synthesis was quantified by measuring the Alexa488 signal per cell using flow cytometry for a population of 500,000 cells, normalizing to signals measured for stained nuclei. Our results revealed a significant reduction in protein synthesis in Δ*pat1* compared to wild-type cells. At 25°C, HPG incorporation in Δ*pat1* cells was reduced to 70.7% of wild-type levels (p = 0.0039, Fig. 5A). This defect was dramatically exacerbated at 34°C, where protein synthesis in Δ*pat1* cells dropped to only 17.8% of wild-type levels (p = 0.0017). These findings demonstrate that protein production is severely impaired in the absence of Pat1, particularly at near-restrictive temperatures.

**Figure 5.**
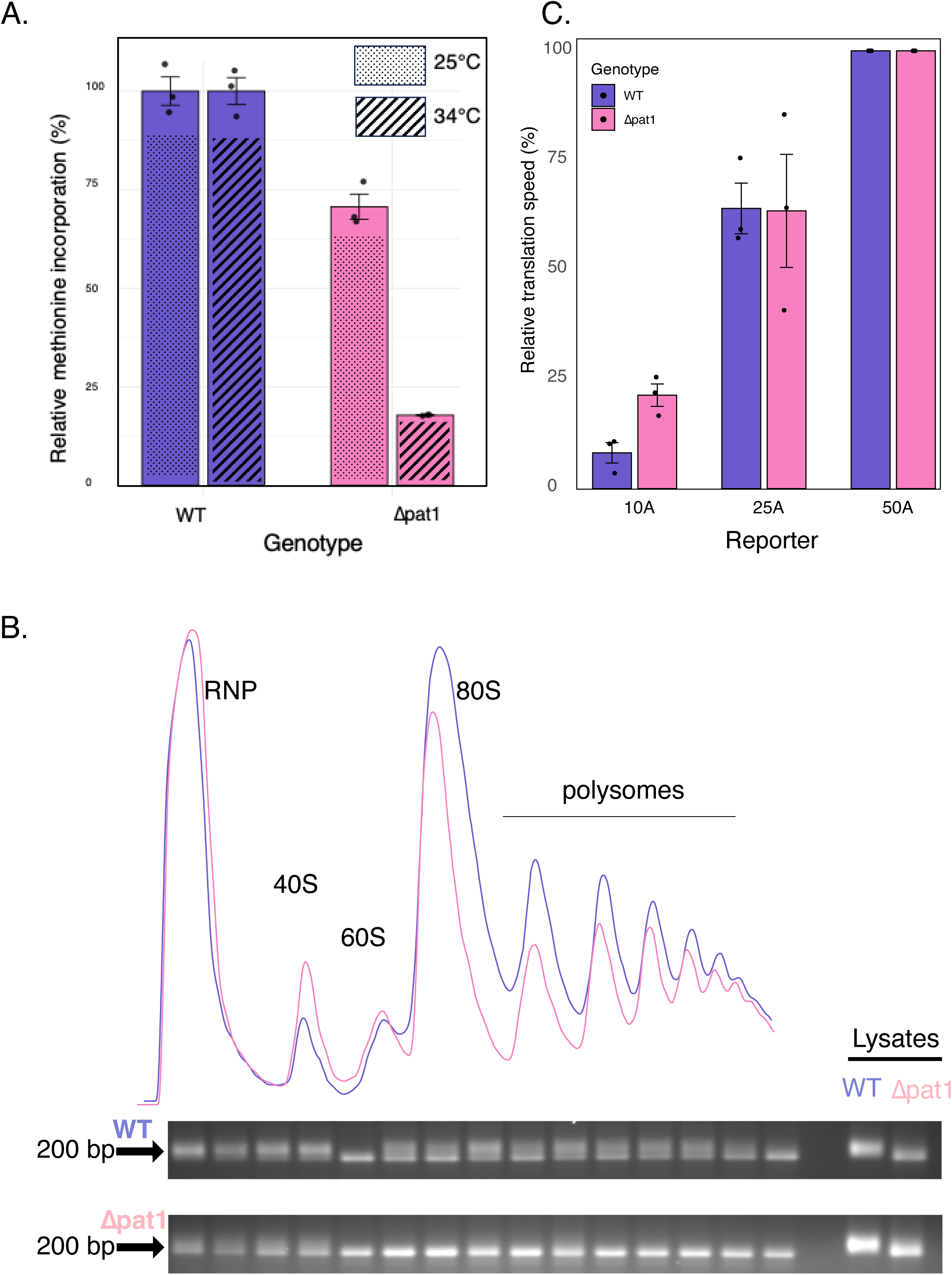
Translation is impaired in Δ*pat1* cells with accumulating short-tailed mRNA being inefficiently translated. (A) Translation efficiency is reduced in Δpat1, a defect exacerbated at 34°C. Translation efficiency was monitored through HPG incorporation for 10 minutes at 25°C or 34°C. Background noise (signal at 0 minutes) was removed before normalization by the nuclear signal for 500 000 cells. The translation efficiency mean of WT triplicates at the cognate temperature was used for normalization (violet) and the translation efficiency of the Δpat1 strain was plotted (pink bars). Individual values (dot) from triplicate biological experiments and standard deviations are presented. t-tests compared WT and Δpat1 mutant at 25°C or 34°C. (B) Oligo(A)-tailed mRNAs are translated in the Δ*pat1* strain. The poly(A) tail size of the RPL41B mRNA was monitored by RACE-PAT across the different polysome fractions and in the original lysate for the wild-type or Δpat1 strains. Polysome profile traces of the wild-type (violet) and Δ*pat1* strain (pink) are shown above ethidium bromide-stained agarose analyses of the RACE- PAT products. RACE-PAT products for of wild-type and Δ*pat1* lysates are shown on the right. Similar results were obtained in 3 biological replicates (Fig. S9). Presence of oligoadenylated mRNA in fraction 5 of the RACE-analysis of wild-type polysomes is unexpected but reproducible in biological replicates (Fig. S9). Mass spectrometry analysis revealed that this fraction contains the CCR4-NOT complex suggesting that deadenylation occurred during polysome fractionation and/or subsequent analyses of the fraction. (C) Oligo(A)-tailed mRNAs are inefficiently translated in both the wild-type and Δ*pat1* strains. Relative translation rates (calculated over 40 minutes) for the wild-type (violet) and Δ*pat1* strain (pink) are derived from triplicate time course experiments (Fig. S10) after transfection of mRNAs encoding NanoLuciferase with different poly(A) tail lengths. Data are normalized in each strain with the NanoLuciferase reporter with a poly(A)-tail length of 50A. Because mRNA transfection efficiencies differ for the wild-type and mutant strains it is not possible to compare the absolute values. The slightly stronger reduction observed for the 10A reporter in the wild-type compared to the Δ*pat1* strain is attributed to exacerbated competition with endogenous mRNAs that have overall longer poly(A) tails in the former case.

Given the high abundance of oligoadenylated mRNAs in Δ*pat1* cells, we hypothesized that reduced translation could either reflect the selective translation of the few mRNAs retaining normal poly(A) tails or, conversely, the inefficient translation of numerous short-tailed mRNAs. To determine whether oligo(A)-tailed mRNAs are translated at 34°C in Δ*pat1* cells, we analyzed polysome profiles and assessed the poly(A) tail length of the RPL41B mRNA using RACE-PAT across the gradient fractions. In wild-type cells, we observed the classical polysome profile, with distinct peaks corresponding to small RNPs, 40S and 60S ribosomal subunits, monosomes, and polysomes. The distribution of RPL41B mRNA poly(A) tail lengths was largely homogeneous across the gradient (Fig. 5B and S9). Long poly(A) tails, predominant in wild-type cells, were distributed across monosome and polysome fractions, with only a slight shift toward shorter tails in heavier fractions. The polysome profile of the Δ*pat1* strain was similar to that of wild-type, albeit with a general reduction in monosome and polysome levels and a slight excess of free 40S subunits, consistent with previous observations^30^. Notably, oligo(A)-tailed mRNAs, which constitute the majority of the transcriptome in Δ*pat1* cells, were distributed across the gradient, indicating that these transcripts are actively translated. The small fraction of mRNAs with longer poly(A) tails was enriched in lighter fractions, likely representing nuclear mRNAs that have not yet been exported to the cytoplasm and thus remained unexposed to deadenylation. These results confirm that oligo(A)-tailed mRNAs in Δ*pat1* cells are indeed translated.

To quantify the impact of poly(A) tail length on translation in wild-type and Δ*PAT1* cells, we constructed NanoLuc reporters with poly(A) tails of 10, 25, or 50 adenine residues (NanoLuc-10A, NanoLuc-25A, and NanoLuc-50A, respectively). Following electroporation of *in vitro*-transcribed, capped mRNAs into wild-type and Δ*pat1* cells^63^, NanoLuc protein levels were measured by luminescence at multiple time points over two hours. The initial rate of protein production was determined for each reporter (Fig. S10). As expected, NanoLuc-50A was the most efficiently translated reporter in both strains and was used to normalize the relative translation rates of the shorter-tailed reporters (Fig. 5C). Reducing the poly(A) tail from 50 to 25 residues had a comparable effect in both wild-type and Δ*pat1* strains, decreasing the initial synthesis rate to 67% and 63%, respectively. Further shortening the tail to 10 residues severely impaired NanoLuc production, reducing the rate to 8% in wild-type and 21% in Δ*pat1* cells, relative to NanoLuc-50A.

Overall, our data indicate that Δ*pat1* cells retain the ability to discriminate against transcripts with short oligo(A) tails, conserving a clear preference for translating mRNAs with longer poly(A) tails. Since the majority of the transcriptome in Δ*pat1* cells consists of short oligo(A)-tailed mRNAs—and these cannot be efficiently degraded due to impaired decapping—this bias negatively impacts the overall translation efficiency especially at near- restrictive temperature.

## Discussion

Efficient cellular function requires mRNA degradation to be as tightly regulated as transcription, with which it is coordinated through transcript buffering. While mRNA poly(A) tail shortening beyond a specific threshold allows a gradual—and in some cells reversible— reduction in protein production, decapping represents an irreversible commitment point in the 5’–3’ mRNA decay pathway, permanently excluding transcripts from the pool of translatable templates. It is therefore critical that the activity of the Dcp2–Dcp1 decapping complex is finely tuned by co-activators. In this context, our data provide compelling evidence for the existence of a general deadenylation-dependent decapping pathway in which Pat1, likely with the Lsm1–7 complex, plays a central role.

Previous studies, largely based on short-read sequencing comparing relative mRNA levels between mutant and wild-type cells, suggested that decapping co-factors, including Pat1, regulate distinct mRNA subsets^28,29,64^. However, these observations were puzzling, as minimal transcriptome alterations produced heterogeneous phenotypes. For example, *PAT1* deletion in yeast results in a strong thermosensitive growth defect, yet this phenotype is associated with only modest changes in gene expression under permissive conditions. Our transcriptome and proteome analyses of wild-type and *Δpat1* cells at near-restrictive temperatures revealed limited, stress-related changes with no bias in protein synthesis. Thus, transcriptome and proteome differences at various temperatures did not explain the poor correlation between growth phenotype severity and modest gene expression changes.

This discrepancy prompted us to examine the transcriptome of these strains using Nanopore-based FLEP-seq2^38–40^. While this method agreed quantitatively with Illumina sequencing (Fig. S2) it uncovered a critical, previously undetected, qualitative difference: a dramatic accumulation of oligo(A)-tailed mRNAs in Δ*pat1* cells. This finding demonstrates that Pat1 affects the entire transcriptome—not just a mRNA subset as previously suggested—and implies that earlier interpretations of the roles of other decapping regulators may also need reevaluation. The near-complete conversion of the *Δpat1* transcriptome to oligo(A)-tailed species reveals that cells lack a homeostatic buffering mechanism that senses poly(A) abundance to maintain it constant; instead, they preserve a constant quantity of transcript. This observation aligns with previous suggestions that Xrn1 participates in buffering, although the precise nature of this mechanism remains unclear^65^.

The capped oligo(A)-tailed mRNAs accumulating in *Δpat1* cells are not mutant-specific species; they are present in wild-type cells with fully functional mRNA degradation pathways (Fig. 4) from which they can be preferentially precipitated with Pat1. This aligns with earlier biochemical analyses showing Pat1’s association with the Lsm1–7 complex, which was reported to bind *in vitro* to 3’ oligo(A)^57^. Our results extend this to an *in vivo* transcriptome-wide property, indicating that deadenylation impacts most cellular mRNAs. Furthermore, the partial suppression of the Δpat1 growth defect by Dcp2/Dcp1 overexpression provides strong evidence for the existence of an important and general deadenylation-dependent decapping pathway in yeast. Structural evidence that Pat1 directly binds Dcp2—and observation that disruption of this interaction impairs decapping *in vivo*—establishes a molecular and functional connection from the oligo(A) tail to cap cleavage^18^. However, since Pat1 also associates with partially deadenylated mRNAs (Fig.4), we cannot exclude that it can, with the associated Lsm1-7, also promote decapping of such species. The role of deadenylation in triggering mRNA decay is widely supported by studies demonstrating how specific factors recruit the main deadenylase to stimulate mRNA turnover^8^. In yeast, Xrn1 also interact with Pat1 suggesting that beyond decapping, deadenylation will also trigger enhanced exonucleolytic degradation of the mRNA body^18^. It is possible that these functions are assumed by Edc4 in mammals^6^.

Recent studies have questioned the existence or importance of deadenylation-dependent decapping^12,13,25^. However, these relied on oligo(dT) enrichment and/or priming that do not allow the detection of mRNAs with short oligo(A)-tails revealed by our work. Additionally, oligo(dT) - based mRNA enrichment strategies, reverse-transcription priming, or sequencing adapter addition may introduce biases in poly(A) tail distribution assessments. These limitations are overcome at the transcriptome level by the FLEP-Seq2 approach, or at the individual mRNA level by RACE-PAT analysis, as they capture only the natural 3’ ends of mRNAs, including the eventual non-A residues. Our findings thus provide robust evidence for a Pat1-driven deadenylation-dependent decapping pathway, potentially overlooked in prior work in part due to technical constraints. The severe growth phenotypes resulting from Pat1 (or Lsm1) inactivation, compared to milder effects resulting from inactivation of other decapping activators like Edc3, further supports the notion that deadenylation-dependent decapping is a global pathway with broad transcriptomic impact. Nevertheless, alternative decapping pathways clearly operate in cells^20,21,23,24,66^ and further studies are needed to quantify their relative contributions to overall decapping under various conditions.

A recent study proposed that yeast Upf1, the central NMD factor, contributes to the decay of mRNAs with shorter poly(A) tails^25^. The short poly(A) tails in that study peaked at ∼20A and are distinct from the oligo(A)-tailed mRNAs identified here. In any case, oligo(A)- tailed mRNAs accumulate massively in Δ*pat1* cells, despite the presence of wild-type Upf1, arguing that Upf1 plays at most a modest role in their degradation. Consistently, Upf1 inactivation in yeast produces no major growth phenotype, suggesting a limited quantitative contribution to overall mRNA turnover. Additionally, the cytoplasmic exosome appears unable to degrade the oligo(A)-tailed mRNAs present in Δ*pat1* cells, possibly due to 3’ ends protection by Lsm proteins and/or insufficient exosome activity.

The high abundance of oligo(A)-tailed mRNAs in the Δ*pat1* strain raised the question of their translational status. Polysome profiling revealed that oligo(A)-tailed mRNAs are translated in Δ*pat1* cells, yet global translation, as monitored by the incorporation of tracer into newly synthesized proteins, is reduced. This defect is exacerbated at near-restrictive temperatures, suggesting that the contribution of the poly(A) tail to protein production is particularly important under suboptimal or stress conditions. Collectively, these observations indicate that inefficient translation of oligo(A)-tailed mRNAs substantially contributes to the thermosensitive growth phenotype of the Δ*pat1* strain. Specifically, reduced translation would lower the rate of cell mass accumulation, matching the slower growth rate of the mutant to approximately maintain overall cellular composition. Supporting this, the transcriptome of the Δ*pat1* strain mimics a stress response, potentially participating in this regulatory adjustment.

Poly(A) tails stimulate translation at the initiation stage by recruiting poly(A)-binding proteins, which interact with eIF4G to facilitate 40S ribosomal subunit loading onto mRNA. However, the translational advantage conferred by poly(A) tails varies across cell types: in some, long tails enhance translation, while in others, tail-length has limited impact on translation or even shorter-tailed transcripts may be more efficiently translated^51,52^. Crucially, all such transcripts retain sufficiently long poly(A) tails to efficiently bind cytoplasmic Poly(A)- Binding Proteins, unlike the oligo(A)-tailed mRNAs accumulating in *Δpat1* cells (although we cannot exclude that in some case this factor could straddle the junction between the oligo(A) tail and the 3’ untranslated region). To test directly whether oligo(A)-tailed mRNAs are translated, we introduced reporter mRNAs with varying tail lengths into cells and monitored protein production. This revealed that mRNAs with oligo(A) tails as short as 10 residues are translated *in vivo*, in both wild-type and *Δpat1* cells, albeit with very low efficiency. This argues that the primary defect in Δ*pat1* cells stems from the poor translation of a transcriptome dominated by oligo(A)-tailed transcripts. Notably, translation efficiency in Δ*pat1* and control cells remains correlated with poly(A) tail length, indicating that no compensatory mechanisms are activated in the mutant to offset the absence of long poly(A) tails.

In summary, our analysis establishes deadenylation-dependent mRNA decay as a transcriptome-wide contributor to the 5’–3’ decay pathway, with Pat1 playing a central role. Thus, Pat1 and its associated Lsm proteins emerge as critical guardians that protect cells from the detrimental effects of poorly translated oligo(A)-tailed mRNAs.

## Methods

### Yeast strains

Yeast strains used in this study are listed in Supplementary Table S1. Gene deletions or TAP-tagging fusions were generated in haploid cells with a one-step PCR replacement protocol using standard lithium acetate transformation protocol^33^. Correct gene tagging or deletion was verified by PCR across both recombination sites using oligonucleotides listed in Supplementary Table S2. For transient overexpression of proteins in specific genetic backgrounds, the same transformation protocol was used to introduce appropriate plasmids (listed in Supplementary Table S3, with details for new constructions described in Supplementary Information).

### Yeast cell culture conditions

Except otherwise stated, for liquid cultures, yeast cells were grown in YPDA (1% yeast extract, 2% peptone, 2% glucose supplemented with 10 mg/L adenine hemisulfate) at 25°C or 34°C and 170 rpm. Prior to HPG labelling, YPDA-grown cells were washed and incubated for 1 hour either at 25°C or 34°C in CSM-Met media (0.67% Yeast Nitrogen Base without amino acids (Becton Dickinson), 2% glucose, amino acids without methionine (MP Biomedicals)). For overexpression of proteins, yeast cells were grown in CSM-Leu-Ura selective media (0.67% Yeast Nitrogen Base without amino acids (Becton Dickinson), 2% glucose, amino acids without leucine and uracil (MP Biomedicals)) prior to monitoring growth in a Mithras LB940 plate reader.

### Total RNA extraction

12 OD_600nm_ of yeast cells in log phase were pelleted and flash-frozen. To extract total RNAs, 0.5 mL of 65°C acidic phenol saturated in water pH 4.0 (Eurobio, CGEXPHE01) were added to frozen pellets. Then were added 0.5 mL of 50 mM NaOAc pH 5.5, 50 µL of 10% SDS, and 0.5 mL of acid-washed glass beads (0.5 mm; BioSpec Products 11079105). Samples were vortexed for 1 minute and incubated for 2 minutes at 65°C before alternating 3 rounds of 30 seconds vortex and 30 seconds incubation at 65°C. After a 10-minute centrifugation at 12 000g, the RNA-containing aqueous phase was extracted twice by the addition of one volume of Phenol:Chlorophrom:Isoamylalcool (25:24:1) (Sigma, Cat# 77617) and subjected to 3 rounds of 30 seconds vortex and 30 seconds incubation at 4°C followed by a 5-minute centrifugation at 12 000g. After centrifugation, total RNAs were precipitated with 1/10^th^ volume of 3M NaOAc and 2 volumes of 100% ethanol. RNAs were washed with 0.5 mL of 70% ethanol, air dried and resuspended in 0.1 mL of autoclaved milli Q water.

### Illumina-based library preparation, sequencing and data processing

Quantities and qualities of total RNAs, as well as final libraries, were evaluated by Qubit fluorometer (Invitrogen^TM^) and Agilent Bioanalyzer (Agilent Technologies). Non-ribosomal RNA libraries were made using the “Library prep for standard total RNA-seq Ribozero/standard quantity” (Truseq^TM^) kit according to the manufacturer’s instructions and sequenced in 50- length Single-Read on a HiSeq 4000 machine (Illumina). Reads resulting from the sequencing were mapped onto the S288C Saccharomyces cerevisiae genome (R64-1-1) with HISAT2 (v 2.2.1)^34^. Mapped reads were assigned and counted with htseq (v 0.13.5)^35^. Differential gene expression analysis was performed on R (v 4.3.2) using DESeq2^36^. Gene ontology analysis were performed on R (v 4.3.2) using clusterProfiler^37^ with annotations from the *Saccharomyces* Genome Database (SGD) using a P-value cutoff of 0.05.

### FLEP-Seq2 library preparation and nanopore sequencing

FLEP-Seq2 libraries were prepared from 3 µg of total RNAs as described in Peter *et al*. 2026^38^, adapted from Long *et al.*, 2021^39^ and Jia *et al.*, 2022^40^ with minor modifications. Briefly, total RNA from *S. cerevisiae* was ribo-depleted using the *Saccharomyces cerevisiae* riboPool kit (siTOOLSs BIOTECH, dp-K096-5) and ligated to 50 pmol of a 5ʹ-riboadenylated DNA oligonucleotide (Universal miRNA Cloning Linker, NEB) in a 20 µl reaction containing 20 U T4 RNA Ligase 2 truncated KQ (NEB, M0373S), 1× T4 RNA Ligase Reaction Buffer, 25% PEG8000, and 40 U RNaseOUT (Life Technologies, 10777019). Ligation was performed at 16°C for 10 h, followed by RNA purification using the RNA Clean & Concentrator-5 kit (ZYMO, R1013). For reverse transcription, ligated RNA was annealed with 2 pmol custom reverse transcription primer (FLEP_seq2_RT) and 10 nmol dNTPs at 65°C for 5 min, then cooled on ice for 2 min. cDNA synthesis was carried out in a 20 µl reaction containing 200 U Maxima H Minus Reverse Transcriptase, Maxima RT Buffer (Thermo Fisher, EP0752), 40 U RNaseOUT, and 20 pmol strand-switching primer (SSPII; ONT, SQK-PCB111.24). Reactions were incubated at 42°C for 90 min, followed by 10 cycles of 50°C for 2 min and 42°C for 2 min, and a final incubation at 85°C for 5 min. cDNA were PCR amplified for 16 cycles in a 25 µl reaction containing 1× PrimeSTAR GXL Buffer (TaKaRa, R050A), 5 nmol of each dNTP, 7.5 pmol of each barcoded primer (Table S2), and 0.625 U PrimeSTAR GXL DNA polymerase (TaKaRa, R050A). For the libraries analyzed by Oxford Nanopore R9, we used the PCR-cDNA Barcoding kit ONT SQK-PCB111.24. For the libraries analyzed by Oxford Nanopore R10, we used ONT SQK-LSK114 and Nanopore barcoded primers (Supplemental Table 2). PCR conditions were as follows: 30s at 98°C; 14 cycles of 10s at 98°C, 15s at 65°C and 10 min at 68°C; finally, 5 min at 68°C. PCR products were incubated for 15 min at 37°C with 20 U of Exonuclease I (NEB, M0293), and denatured for 15 min at 80°C. For each sample, 6 PCR reactions were pooled. PCR products were cleaned-up with 3 successive rounds of purifications with the NucleoMag NGS Clean-up and Size Select Kit (Machery-Nagel, 744970.50), using bead ratio of 0.7X, 0.55X and 0.55X for rounds 1, 2 and 3, respectively. Libraries quantities and qualities were verified as described previously for the Illumina-based library preparation. Libraries were sequenced for ∼72 hours on a P2 PromethION 2 Solo device (Oxford Nanopore) using a FLO-PRO002 (R9.4.1) or FLO-PRO114M (R10. 4.1).

### Computational analysis for FLEP-Seq2

The raw signal FAST5 files generated by Oxford Nanopore R9 were base-called using Guppy (Version 6.3.8+d9e0f64) with default parameters using the high accuracy model ((-c dna_r9.4.1_450bps_hac). Reads with a mean quality score inferior to 7 were filtered out. Reads were mapped to the *S. cerevisiae* S288C genome (R64-1-1) using Minimap2 (2.26- r1175). Transcript 3’ end detection and poly(A)-tail length determination were performed using a pipeline adapted from^40^ and available on Github (Peter et al. 2026)^38^.

The raw signal POD5 files generated by Oxford Nanopore R10 were base-called using Dorado v0.9.0 (model: dna_r10.4.1_e8.2_400bps_sup@v4.3.0) using poly(A) tail estimation (-- estimate-poly-a) and adapter sequences specified in the polya_config.toml file (front_primer = TTTCTGTTGGTGCTGATATTGCT; rear_primer = ATTGATGGTGCCTACAG). Reads were aligned to the *S. cerevisiae* S288C genome (R64-1-1) using minimap2 implemented in Dorado (--mm2- opts “-x splice”), and only primary alignments were retained. Read start and end coordinates, estimated poly(A) tail lengths, annotations, and right soft-clipped sequences were extracted from the aligned BAM files using a custom Python 3 script. The soft-clipped sequence corresponding to the 3’ end was further parsed into the poly(A) tail and 3ʹ adapter.

Only nuclear-encoded mRNAs, based on the S288C annotation, were used for analyses. To analyze the distribution of poly(A)-tail lengths individual mRNA molecules were grouped by gene after removing rare outlier transcripts with poly(A) tails longer than 100 residues. To avoid biases due to highly abundant or rare transcripts, the poly(A) tail profile was generated by summing the relative distributions for every gene with at least 25 mRNA molecules analyzed. For heatmaps, the same filters were used and poly(A)-tails size distribution in 10- nucleotide bins from 0 to 100 adenosines residues were plotted after z-score rescaling.

For the analysis of transcripts bound by Pat1 or Pab1, the percentage of transcripts with less than 10 adenosines was calculated from the whole transcript counts while the poly(A) tail profile between 10 and 100 residues was generated by summing the relative distributions for every gene with at least 25 mRNA molecules detected.

### Immuno-precipitation of protein-bound mRNAs

40 OD_600nm_ of exponentially growing yeast cells were harvested by a 5 minutes centrifugation at 4 500 g, washed in Tris-Cl pH 8.0 and resuspended in 0.4 mL of IPP150 (10 mM Tris-Cl pH 8.0; 150 mM NaCl; 1X Protease Inhibitor Cocktail without EDTA (Roche) and 0.1% IGEPAL®CA-630 (Sigma-Aldrich)) before cell lysis with 0.4 mL of acid-washed glass beads (0.5 mm; BioSpec Products 11079105) with a FastPrep®-24 Classic bead beating grinder and lysis system (MP Biomedical) for 40 seconds at 6 m/s. The liquid phase was recovered, and subjected to two rounds of centrifugation at 16 000 g at 4°C for 5 minutes. 1 volume of the supernatant was supplemented with 40 U of RNAse inhibitor SUPERaseIn^TM^ (ThermoFisher) as well as ¼ volume of IPP150-washed Protein G Dynabeads^TM^ (ThermoFisher) and incubated for 1 hour at 4°C on a slow-spinning wheel (5 rpm). Beads were washed 4 times with IPP150 without Protease Inhibitor Cocktail and RNA elution was performed for 45 minutes at 50°C through the addition of 100 µg of Proteinase K in 100 mM Tris-HCl pH 7.5, 12.5 mM EDTA pH 8.0, 150 mM NaCl, 1% SDS. RNAs were extracted through two rounds of Phenol:Chloroform:Isoamylalcool (25:24:1) purification and ethanol precipitated.

### Immunoprecipitation of capped mRNAs

5 mg of IPPH-washed (500 mm NaCl; 10 mM Tris-Cl pH 7.5; 0.1% IGEPAL®CA-630 (Sigma-Aldrich)) Protein G Dynabeads^TM^ (ThermoFisher) were resuspended in 1 mL IPPH supplemented with 0.1 U/µL of RNAse inhibitor SUPERaseIn^TM^ (ThermoFisher) and with 10 µg/mL of Anti-7-methylguanosine (m7G) monoclonal Antibody RN017M (MBL), or as control 10 µg/mL of peroxidase anti-peroxidase (Sigma P1291) and incubated overnight at 4°C, 5 rpm. Beads were washed 5 times with 0.5 mL IPPL buffer (150 mM NaCl; 10 mM Tris-Cl pH 7.5; 1 mM EDTA pH 8; 0.05% IGEPAL®CA-630 (Sigma-Aldrich)). Beads were resuspended in 0.2 mL IPPL buffer. 50 µg of total RNA were incorporated to half a volume of IPPL-resuspended beads and 40 U RNAse inhibitor SUPERaseIn^TM^ (ThermoFisher) were added. After 8 hours of incubation on a slow-spinning wheel (5 rpm) at 4°C, beads with bound mRNAs were recovered with the help of a magnet and resuspended in 90 µL of IPPL and conserved at 4°C. In parallel, the first supernatant was added to the other half of IPPL-resuspended beads and incubated overnight at 4°C, 5 rpm to recover any remaining capped-RNA. The following morning, the second supernatant was discarded and the first and second bead pellets were resuspended together and adjusted to 1% SDS. Beads were incubated at RT for 1 hour at 15 rpm and eluted RNAs were extracted through two rounds of Phenol:Chloroform:Isoamylalcool (25:24:1) purification and ethanol precipitated.

### Assaying the sensitivity of mRNAs to degradation by Xrn1

For biological triplicates, identical quantities of total yeast mRNAs were mock-treated, treated with Xrn1 (NEB M0338), or first treated with the RppH (NEB M0356S) enzyme to remove mRNA caps before Xrn1 treatment, essentially as recommended by the manufacturer. For each sample, RT-qPCR reactions for different test genes were performed in technical triplicates. The differences of the (technical) average of the Ct obtained for the different samples with the value obtained for the decapped and Xrn1 treated sample (theoretically most susceptible to RNA degradation) were computed per gene. The corresponding fold differences, reflecting “sensitivity” to Xrn1 degradation, were plotted after averaging for biological triplicates.

### Lysate preparation for polysome and LC-MS/MS analyses

Exponentially growing cells (100mL at 0.8 OD_600nm_) were supplemented with cycloheximide to a final concentration of 100 µg/mL. Cells were washed 2 times with polysome lysis buffer (PLB) (10 mM Tris-Cl pH 7.5; 100 mM KCl; 5 mM MgCl_2_; 3 mM DTT; 100 µg/mL cycloheximide) and resuspended in 0.5 mL PLB. 0.5 mL of acid-washed glass beads (0.5 mm; BioSpec Products, 11079105) were added and cells were lysed at 4°C by alternating 1-minute vortexing and 1-minute ice incubation. The supernatant was centrifuged twice at 20 000g for 10 minutes at 4°C in an Eppendorf 5417 C centrifuge before polysome- or LC-MS/MS -analyses. Procedures for analyses by mass spectrometry are detailed in Supplementary Information.

### Polysome analyses

Polysome fractionation was made on 10-45% sucrose gradients in PLB 1X. Gradients were formed by introducing 6 mL of 10% sucrose-PLB 1X at the bottom of a polypropylene “Beckman Coulter” tube (14x89 mm), followed by 6 mL of 45% sucrose-PLB 1X, placed beneath the 10%-sucrose PLB 1X with the help of a syringe. Tubes were closed with 4-mm caps and linear gradients generated with the “Gradient Master Biocomp” (Biocomp Instruments) with the “Short, sucrose gradient 10-45%” program. 10 OD_260nm_ from the yeast lysates were layered drop-wise on the gradients before centrifugation in an SW41 rotor for 2 hours at 40 000 rpm at 4°C in a Beckman Coulter Optmima LE-80K Ultracentrifuge. The absorbance profile at 254 nm was recorded through the ISCO Teledyne Foxy Jr apparatus and fractions of about 600 µL were recovered.

### RACE-PAT analyses

RACE-PAT analyses were made as described in Mauxion and Séraphin, 2024^41^. Briefly, up to 1 µg of total or immunoprecipitated RNAs were ligated to 50 pmol of RNA ligation oligonucleotide (listed in Table S2) in a 10 µL reaction containing 10 U T4 RNA ligase 1 (NEB), 1X T4 RNA Ligase RNA Buffer (50 mM Tris-HCl pH = 7.5; 10 mM MgCl_2_; 1 mM DTT), 10% PEG 8000 1X and 1 nmol ATP for 2 hours at 25°C before enzyme inactivation 10 minutes at 70°C. Half of the ligated-RNA reaction was mixed with 5 µL of milli Q water and 100 pmol of RT primer, incubated for 5 minutes at 65°C and at least 2 minutes at 4°C. Reverse transcription mix containing 1X Transcriptase inverse RevertAid H Minus Buffer 1X (ThermoFisher), 20 U of RNAse inhibitor SUPERaseIn^TM^ (ThermoFisher) and 200 U of Transcriptase inverse RevertAid H Minus were added to a final volume of 20 µL and incubated for 1 hour at 42°C before enzyme inactivation for 10 minutes at 70°C. 1 µL of the resulting cDNA was used for 3’ end amplification by the DreamTaq DNA polymerase (ThermoFisher) according to the manufacturer’s instructions with a maximum of 25 cycles. Primers are listed in Table S2.

### Monitoring translation efficiency by HPG incorporation

Translation efficiency assays were performed as in Basier and Nurse, 2023^42^ with minor changes. Briefly, 20 mL of exponentially grown yeast cells at OD_600nm_ = 0.3 were washed and resuspended in 20 mL of CSM-Met media. After incubation for 1 hour at 25°C or 34°C, HPG was added to a final 10 µM concentration from a 50 mM stock. A first sample of 3.84 mL of cells was directly fixed with 1.16 mL of 16% methanol free paraformaldehyde (Electron Microscopy Sciences), vortexed for 5 seconds and incubated at room temperature in the dark on a rocker for 40 minutes. A second sample was harvested after 10 minutes incubation with HPG and fixed the same way, and incubated in the dark on a rocker for 30 minutes. Cells from both samples were spun at 2000g for 5 minutes and washed in Phosphate Buffer Saline (PBS) 1X supplemented with 1% (w/v) BSA, then resuspended in PBS 1X, 1% (w/v) BSA, 1% (v/v) Triton X-100 and incubated for 30 minutes in the dark on a rocker. Cells were resuspended in PBS 1X with 1% (w/v) BSA and incubated for 1 hour in the dark on a rocker. To label HPG the Click-iT^TM^ HPG Alexa Fluor^TM^ 594 Protein Synthesis Assay Kit (ThermoFisher) was used. Cells were washed with 500 µL 1X Click-iT reaction buffer, resuspended in 500 µL of Click-iT reaction cocktail and incubated for 30 minutes in the dark at 1000 rpm. Cells were spun for 15 seconds at 17 000g and washed in 50 mM sodium citrate, before a 10-minute incubation with 50 mM sodium citrate supplemented with 1X Nuclear-ID blue (ThermoFisher). Cells were washed in 5 mM sodium citrate before being resuspended in 500 µL of the same buffer and kept at 4°C for 1 hour in the dark before flow cytometry analysis (see Supplementary data).

### Spheroplast preparation for mRNA electroporation

Spheroplasts were prepared as in Taverniti and Séraphin, 2015^43^ with minor changes. 15 mL of exponentially growing cells at 0.8 OD_600nm_ were spun 5 minutes at 4 500g at RT and resuspended in 5 mL buffer A (1M sorbitol, 15 mM β-mercaptoethanol, 50 mM Tris-HCl pH 7.5, 1 mM MgCl_2_, 30 mM DTT) before addition of 1 mL Zymolase (1 mg/mL). Cells were incubated for 30 minutes in a slow-spinning wheel, centrifugated 5 minutes at 500 g at room temperature and washed twice in 5 mL of buffer A. Digested cells were resuspended in 1 mL buffer A and added to 10 mL YPDA-sorbitol 1M. Spheroplasts were left to regenerate for 90 minutes at 25°C then washed twice in 10 mL 1M sorbitol. Cells were resuspended to 5 OD_600nm_ and 180 µL were aliquoted in a tube for subsequent electroporation.

### In vitro transcription and mRNA capping, purification and electroporation

Plasmids encoding the firefly mRNA were digested with 25U of BsqBI restriction enzyme (NEB) and purified by Phenol:Cholorform:Isoamylalcool (25:24:1) and precipitation before being transcribed *in vitro* with the Hiscribe T7 High Yield RNA synthesis kit (NEB) from 1 µg of template DNA following the manufacturer’s instructions.

After transcription, 2 µL of DNAse I (Thermo Scientific) were added and the reaction was incubated for 15 minutes at 37°C. mRNA production, integrity and size were verified on a 2% agarose gel with 0.5 µL of the reaction.

100 µg of the resulting mRNAs were capped with the Faustovirus Capping Enzyme (NEB) according to the manufacturer’s instructions. Capped mRNAs were purified with the Monarch Spin RNA Cleanup Kit (500 µg) (NEB) according to the manufacturer’s instructions and kept at −80°C until further use. 1 µg of the purified capped mRNA was mixed to 180 µL spheroplasts and placed on ice. The mix was transferred to a MicroPulser electroporation cuvette with a 0.2 cm gap (Bio-Rad) and electroporated on a BioRad GenePulser system with settings 800V, 25 µF, 1000 Ohm as previously described^44^.

### NanoLuc Translation rate assessment

The totality of electroporated spheroplasts were transferred to a round-bottom 14 mL Falcon tube and supplemented with 500 µL of YPDA with 1M sorbitol and incubated at 25°C or 34°C and 70 rpm. For NanoLuciferase measurements, at each time point (0, 5, 10, 20, 40, 60 and 120 minutes), 50 µL of the cultures spheroplasts were added to 50 µL of Nano-Glo® Luciferase Assay reagent (Promega) (1 volume of Nano-Glo® Luciferase Assay substrate with 50 volumes of Nano-Glo® Luciferase Assay buffer) in a 5 mL clear tube. The signal was directly measured with the Lumat LB 9507 (Berthold Technologies) apparatus, with a time detection of 2 seconds. Initial translation rate was measured by calculating the linear regression after 40 minutes incubation of each reporter as described previously^44^.

### Statistical analyses

All experiments were performed in biological triplicates, unless stated otherwise. All correlation analyses were performed using Spearman’s rank correlation coefficient. A Wilcoxon test was made to assess if the percentage of transcripts with retained introns were significantly different. HPG incorporation differences were compared with a t-test. Values with a p < 0.05 were deemed statistically significant. Statistical data were calculated by R.

## Acknowledgements

We thank members of our teams and C. Charenton for critical reading of the manuscript and suggestions. We thank the Proteomics Plateform, flow cytometry (CytoEast) and Genomeast Plateform of the IGBMC. We thank the Gene expression plateform of the IBMP and H. Yu for advice in loading the libraries on the Oxford Nanopore flowcell. We would also like to thank Dr. C. Gaudon-Plesse, Dr. A. Dziembowski and Dr. V. Taverniti for plasmid gifts, as well as Dr. V. Roudko and C. Faux for yeast strains.

## Author contributions

Experimental planning and writing of original draft: L.L. and B.S.; experiment realization: L.L., and B.S., data analyses: L.L., J.P., H.Z. and B.S., figure preparation: L.L. and H.Z., manuscript finalization: L.L, H.Z. and B.S., project supervision: B.S., funding: B.S.

## Data availability

NGS datasets generated during this study have been deposited in NCBI’sGene Expression Omnibus and are accessible through GEO Series accession number GSE338293.

Data for Nanopore FLEP-Seq2 have been deposited an SRA with the accession number PRJNA1496879.

Mass spectrometry proteomics raw data have been deposited to the ProteomeXchange Consortium via the PRIDE partner repository with dataset identifiers PXD081268.

## Materials and correspondence

Described material is available upon reasonable request from the corresponding author.

